# Unveiling patterns of roadkill of a migratory amphibian in Hong Kong with implications for mitigation

**DOI:** 10.1101/2025.07.04.663120

**Authors:** Nicolas Louis Michel Brualla, Gena Yip, Evan John Pickett, Hon Shing Fung, Anthony Lau

**Affiliations:** Science Unit, School of Interdisciplinary Studies, Lingnan University, Hong Kong SAR, China; Faculty of Science, The University of Hong Kong, Hong Kong SAR, China; Frigatefilms, Hong Kong SAR, China

## Abstract

Road-related mortality, particularly wildlife-vehicle collisions, poses a significant threat to amphibian populations, with billions of individuals falling victim annually. The rapid urban development in Hong Kong, China, including the forthcoming construction of a “Northern Metropolis” area, is expected to bring over 2.5 million new residents, potentially increasing traffic and posing a heightened threat to local amphibians during their migration periods. To help prevent future biodiversity loss, our study focuses on the near-threatened newt species *Paramesotriton hongkongensis* in Hong Kong, investigating the spatiotemporal distribution of roadkill during migration seasons. Weekly surveys at four hotspots revealed 1,563 animal carcasses, the majority of which were *P. hongkongensis*. Factors triggering mass mortality events were explored, although no significant correlations were found. Prediction models demonstrated moderate accuracy in detecting mass mortality events, indicating a need for further refinement. Recommendations for site-specific mitigation measures to protect amphibians during their migrations are discussed, with an emphasis on the importance of conducting fine-scale surveys for effective conservation strategies.

## Introduction

Roads impose many negative impacts on wildlife, such as direct mortality, interruption of animal movements, and habitat fragmentation, all leading to population isolation and decline (Beebee, 2013; Ochs, Swihart & Saunders, 2024; Pinto et al., 2024). Direct mortality caused by wildlife-vehicle collisions (i.e., roadkill) has the most negative impact on local populations, with billions of animals killed annually (Grilo et al., 2020, 2025). Roadkill numbers of small vertebrates such as amphibians, reptiles, and small mammals are often underestimated, particularly in surveys conducted in moving vehicles, because of their size and low carcass persistence (Glista, Devault & DeWoody, 2008; Rytwinski & Fahrig, 2015; Wang et al., 2022). Still, various studies showed that amphibians can account for up to 90% of total roadkill (i.e., Beebee, 2013; D’Amico et al., 2015; Wang et al., 2022).

Amphibians are especially susceptible to roadkill due to several ecological, physiological, and behavioural characteristics. Many amphibians seasonally migrate between terrestrial and aquatic habitats to complete their biphasic life cycles (Kovar et al., 2009; Semlitsch, 2008; Sinsch, 2014). Also, amphibian activities are highly correlated with meteorological conditions. Humid and warm weather is usually favourable for amphibians to migrate, forage, reproduce, and disperse en masse (Dervo et al., 2016; Todd & Winne, 2006; Ximenez & Tozetti, 2015). In addition, amphibians typically exhibit slow movement and a tendency to freeze when confronted by approaching vehicles (Bouchard et al., 2009; Lima et al., 2015; Mazerolle, Huot, & Gravel, 2005). Therefore, mass roadkill events of amphibians are more likely to happen on days with favourable weather during their migration period (Garriga et al., 2017; Mestre et al., 2019; Zhang et al., 2018).

Various permanent and temporary mitigation measures have been proposed and implemented to alleviate the problem of mass amphibian roadkill. One of the commonly applied measures is crossing tunnels (i.e., culverts) with guiding fences, which reduces animal mortality on roads (Boyle et al., 2021; Helldin & Petrovan, 2019; Pinto et al., 2024). To maximize the effectiveness of these structures, it is necessary to identify roadkill hotspots, where animals concentrate their crossings (Langen et al., 2009; Zhang et al., 2018; Zimmermann Teixeira et al., 2017). However, building these infrastructures is costly and time-consuming (Beebee, 2013). A more immediate mitigation includes short-term road management practices such as traffic detour or temporary road closure (Driessen, 2021; Pokorny et al., 2022; Zhang et al., 2018). These mitigations require investigation of the temporal pattern of roadkill and factors that influence the roadkill number (D’Amico et al., 2015; Raymond et al., 2021).

Studies have shown that meteorological conditions such as temperature, rainfall, and relative humidity are positively related to amphibian roadkill (Garriga et al., 2017; Mestre et al., 2019; Zhao et al., 2023). However, to date, most temporal studies have been conducted on a seasonal or monthly scale, which is too broad for temporary mitigations to be effectively applied on roads of frequent use (Arévalo et al., 2017; Jia et al., 2024; Wang et al., 2022). Roadkill studies examining finer temporal patterns (daily or weekly) in conjunction with meteorological conditions would help predict the exact temporal peaks of amphibian roadkill and inform traffic constraints to be applied accordingly. This way, mitigations allow less disturbance to the public and are more cost-effective.

This study tested the feasibility of predicting fine-scaled temporal peaks of roadkill for a near-threatened newt species *Paramesotriton hongkongensis* (Hong Kong newt). This tropical newt species migrates between their breeding streams and core terrestrial habitats during early spring and autumn (Fu, Karraker & Dudgeon, 2013; Lau et al., 2017). Mass roadkill events during early spring have been observed for years, yet only one pilot study on a residential village road is available, which recorded 40 newt roadkill carcasses from five surveys (Lau, 2017).

Furthermore, ongoing urban development in Hong Kong, China is expected to accommodate over 2.5 million new residents and poses a new challenges for local amphibians (Development Bureau of Hong Kong Special Administrative Region Government, 2021). This development is likely to intensify traffic around the city, raising concerns about increased mortality rates for amphibians during their migratory seasons. This situation underscores the urgent need for proactive conservation measures to mitigate the escalating threats facing the Hong Kong newt and other vulnerable amphibian species in the region.

A clearer understanding of when, where, and what triggers mass roadkill events of these newts is needed to guide mitigation efforts. Thus, the objectives of this study are 1) to investigate the fine-scaled spatial and temporal distribution in four roadkill hotspots of Hong Kong newt during post-breeding migration; 2) to assess the feasibility of predicting mass roadkill event by weather conditions (temperature, rainfall and relative humidity); and 3) to recommend permanent and temporary mitigation to alleviate amphibian roadkill during their migration period.

## Materials & Methods

### Survey area

Hong Kong is a city of more than 7.5 millions inhabitants. It is characterized by subtropical climate with distinct dry and wet seasons regulating the migration cycles of amphibians. Despite being a heavily populated area with rapid urban development, roughly 76% of its land is covered by natural habitats (Development Bureau of Hong Kong Special Administrative Region Government, 2018). We selected four survey sites based on their historical records of newt migration occurrences. All four sites share common elements with a proximity to known newt breeding streams and a paved road with frequent traffic (Figure 1). The first site is a road transect inside Chuen Lung (CL) village, situated at ∼305 m ASL on the west slope of Tai Mo Shan Mountain (the highest peak in Hong Kong continental territories), next to the Chuen Long River (Figure 1). The second survey transect runs across Pak Ngau Shek (PNS) village, located at a lower altitude (∼75 m ASL) on the northern slope of Tai Mo Shan, and with the Lam Tsuen River crossing the transect several times. The third survey site is a paved portion of Mui Tsz Lam (MTL) road leaving Tai Shui Hang and running into MTL village, at the lowest altitude of our sampling (∼45 m ASL) (Figure 1). The Tai Shui Hang stream is located near this transect, and low urban development is present, potentially reducing the traffic volume. Lastly, the fourth site is along Fei Ngo Shan Road, next to Kowloon Peak (KP) mountain, located on the Southern part of Ma On Shan Country Park, with the highest elevation of all four sites (∼325 m ASL) (Figure 1). A stream runs along most of the transect, and the immediate surroundings are less urbanised than the other selected sites. The respective distances for each transect are 900m (CL), 2300 m (PNS), and 1300 m for the last two sites (MTL and KP).

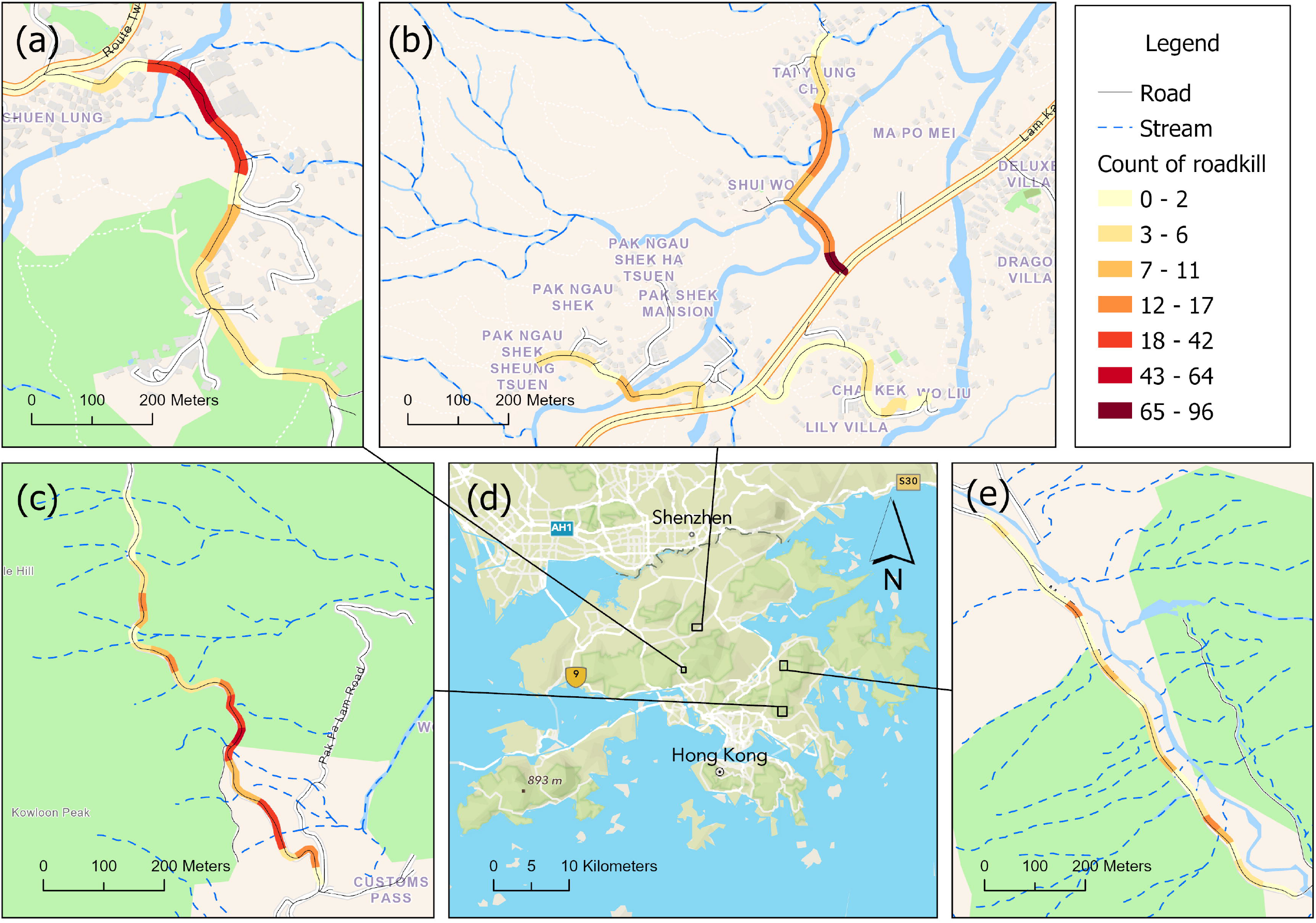

### Data collection

The first objective was to quantify the number of *P. hongkongensis* roadkill at the four study sites. We recruited and trained 20 citizen scientists to conduct field surveys from March 19^th^ to May 28^th^, 2024. Each site was surveyed once a week, with additional *ad hoc* surveys after heavy rain episodes. Surveyors collected roadkill data using a mobile application (TrailWatch app) on their smartphone to report the GPS coordinates of each newt carcass, identifying any non-newt carcasses, taking pictures of the carcasses with a ruler, counting the number of cars seen driving on the survey road during the survey, and lastly, recording the time spent on each survey. Data from the surveys have been organized in a single table for further analysis (Supplementary Table 1).

Ethical approvals were granted by the Hong Kong Department of Health ((22-2) in DH/HT&A/8/2/8 Pt.3) and Lingnan University (EC052/2021). Approval were to collect specimens in Country Parks was granted by the Agriculture, Fisheries, and Conservation Department ((21) in AF GR CON 09/51 Pt.10).

Daily rainfall, relative humidity, and temperature data for the three days prior to each survey were collected from the Hong Kong Observatory Open Data website, using the closest weather station of each survey site: Kowloon Peak – TC 22.342274, 114.227805 // Mui Tsz Lam – SHA 22.395708, 114.230586 // Chuen Lung – TWN(RF)/TW(RH) 22.395385, 114.109803 // Lam Tsuen (PNS) - SEK 22.444013, 114.127581.

### Data analyses

#### Descriptive statistics

Before any analysis, we prepared our main variable representing the roadkill by dividing the number of newt carcasses by the distance of the transects (roadkill per kilometre, RpK) for each survey. We sorted each survey by RpK (high to low) and calculated the cumulative RpK as a proportion of total cumulative RpK, allowing us to define Mass Mortality Events (MMEs, i.e., surveys with the most carcasses and representing 90% of the total of roadkill). We worked on the temporal distribution of RpK to illustrate the expected short seasonality aspect of the newt migrations and to allow discussion on the best mitigation measures’ recommendations. Additionally, we illustrated the spatial distribution of roadkill along the four different sites using a geospatial information system. This step was crucial as decisions on optimal mitigation measures would also change depending on a clustered or random distribution of carcasses on site.

#### Generalized linear models and prediction models

Once we identified the roadkill spatial distribution, we aimed to determine the factors triggering MME during the migratory season, using binomial generalised linear models (GLM). We used five different variables known from previous studies that influence amphibian road mortality. Average temperature (°C), average relative humidity (%) and cumulative rainfall (mm) for the three days prior to the survey date were used as these are common factors triggering amphibian activity (e.g., Pinto et al., 2024). Traffic volume, measured as the number of vehicles per hour during each survey, was also considered, as traffic is a direct contributor to mortality. Lastly, we added a categorical variable “site” to account for any potential variation due to the four sites. We created 15 candidate models comprising combinations of two to four of the above variables, and we ran those models following a binomial distribution. Due to limited sample size, the global model including all five variables could not be run due to over-parameterisation. We identified the factors impacting probability of MME by selecting the best models according to ΔAICc. We averaged the beta estimates from all models weighted according to AICc to obtain a more robust and complete model to use for predictions. Models without a variable were averaged as if the beta estimate were zero. In addition, we looked for the confidence intervals of those averaged beta estimates, allowing us to describe the ascending or decreasing effect of each tested factors.

Our averaged model was tested with different conditions for prediction models. First, we ran a prediction model using the parameters of the averaged model and the original data. Then, we ran a second prediction model based only on weather forecast data and fixing the traffic data constrained to the mean cars counted per hours as we are not able to forecast the traffic in the model for future predictions. In addition, it can be relevant to understand the impact of changing the traffic data on the prediction models, as traffic regulation could be part of the suggested mitigations. Thus, the proportions of correctly assigned MME (and absence of MME) were compared between prediction models and the original surveyed data to determine the accuracy of those models.

## Results

### Roadkill surveys

We counted 1,563 animal carcasses in our 11 weeks survey, a majority of which were *P. hongkongensis* (91.3%; Table 1). We also observed roadkill of 3 species of frogs, 2 species of lizards, 3 species of snakes, 1 bird, and 4 species of invertebrates, but none of them exceeded 10 individuals (Table 1). More than 60% of the newt roadkill were from one site (KP; 870 individuals), with another two sites (CL and PNS) contributing similar numbers (243 and 218 individuals, respectively) (Table 1). The mean number of newts killed per kilometre of road surveyed was 20.76 (SD = 31.67; range =0-136.92), and right-skewed (median = 5.56), where several days at KP and CL have high RpK (Figure 2). Those isolated MMEs represent 37.74% of the survey effort (20 out of 53) and contribute to 90% of the total RpK of newts (Figure 3). The highest numbers of newt carcasses were found at KP, during April (Figure 4). This peak of roadkill is illustrated on all four sites, despite the heterogeneity of the RpK’s distribution observed along the remaining survey period (Supplementary Figure 1). Overall roadkill numbers fluctuated according to the average rainfall values (Figure 4). However, site-specific variations are visible, with KP having the roadkill numbers following weather data variations, and MTL having contrasting results with most carcasses found prior to the major increase in rainfall (Supplementary Figure 1). Spatial distribution along the surveyed roads is also site-specific (Figure 1). Roadkill along KP and MTL roads are evenly distributed (Figure 1c and 1e), contrasting with the clustered distributions observed along PNS and CL roads (Figure 1a and 1b).

**Table 1:** Listing of roadkill per species and per site, illustrating the over-representation of P. hongkongensis on the Hong Kong’s roads.

| Scientific name | Common name | CL | KP | MTL | PNS | Total |
| --- | --- | --- | --- | --- | --- | --- |
| Amphibians |  |  |  |  |  |  |
| *Paramesotriton hongkongensis* | Hong Kong newt | 243 | 870 | 96 | 218 | 1427 |
| *Duttaphrynus melanostictus* | Asian common toad | 2 | 3 | 2 |  | 7 |
| *Hylarana latouchii* | Brown Wood Frog | 1 |  |  |  | 1 |
| *Kaloula pulchra* | Banded bullfrog |  |  |  | 1 | 1 |
|  | Other frogs | 6 | 10 | 5 | 66 | 87 |
| Lizards |  |  |  |  |  |  |
| *Calotes versicolor* | Changeable lizard | 1 | 1 | 3 |  | 5 |
| Scincidae | Skinks |  | 1 |  |  | 1 |
|  | Other lizards | 2 | 1 | 1 | 2 | 6 |
| Snakes |  |  |  |  |  |  |
| *Rhabdophis subminiatus* | Red-necked keelback | 1 |  |  |  | 1 |
| *Trimeresurus stejnegeri* | Bamboo pit viper | 1 |  |  |  | 1 |
| *Ptyas major* | Greater green snake | 1 |  |  |  | 1 |
|  | Other snakes | 4 | 6 |  |  | 10 |
| Birds |  |  |  |  |  |  |
| Aves | Birds |  | 1 |  |  | 1 |
| Others |  |  |  |  |  |  |
| Lumbricina | Earthworms | 3 |  |  |  | 3 |
| Gastropoda | Slugs | 1 |  |  |  | 1 |
| Gastropoda | Snails | 1 |  |  |  | 1 |
| *Gaeana maculata* | Speckled black cicada |  |  | 1 |  | 1 |
| Non-identified |  | 4 | 2 | 4 | 5 | 15 |
| TOTAL |  |  |  |  |  | 1563 |

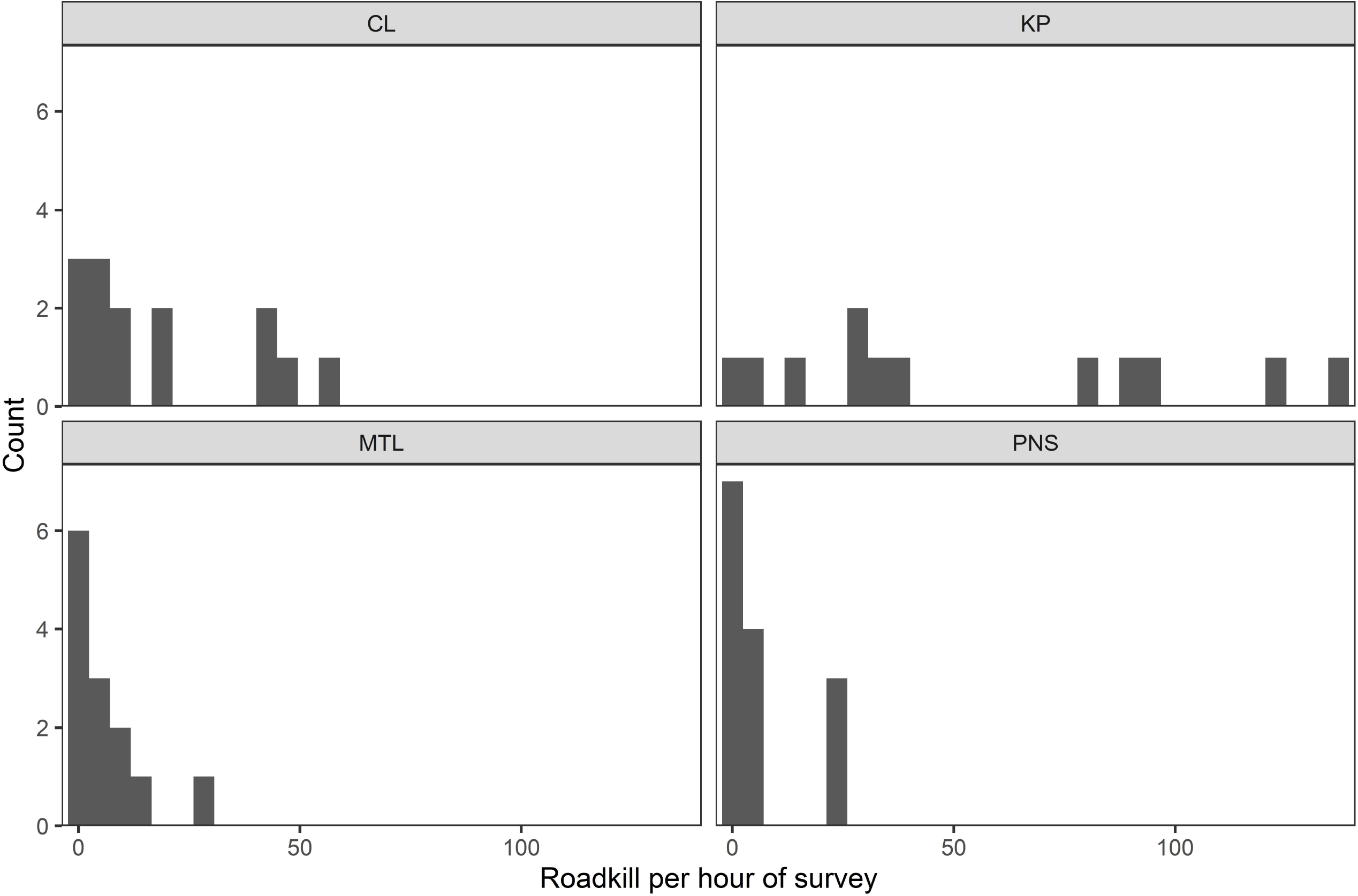

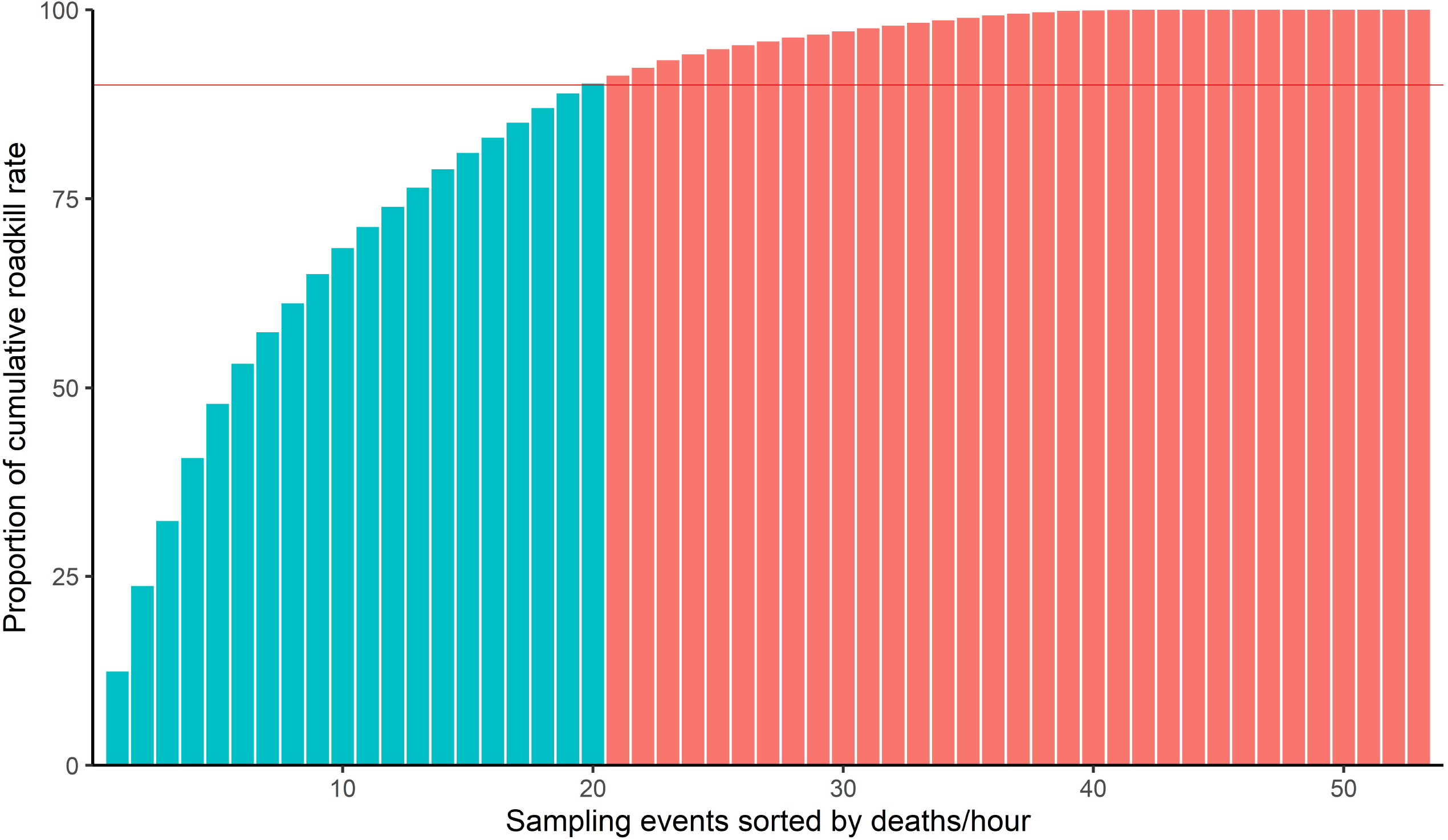

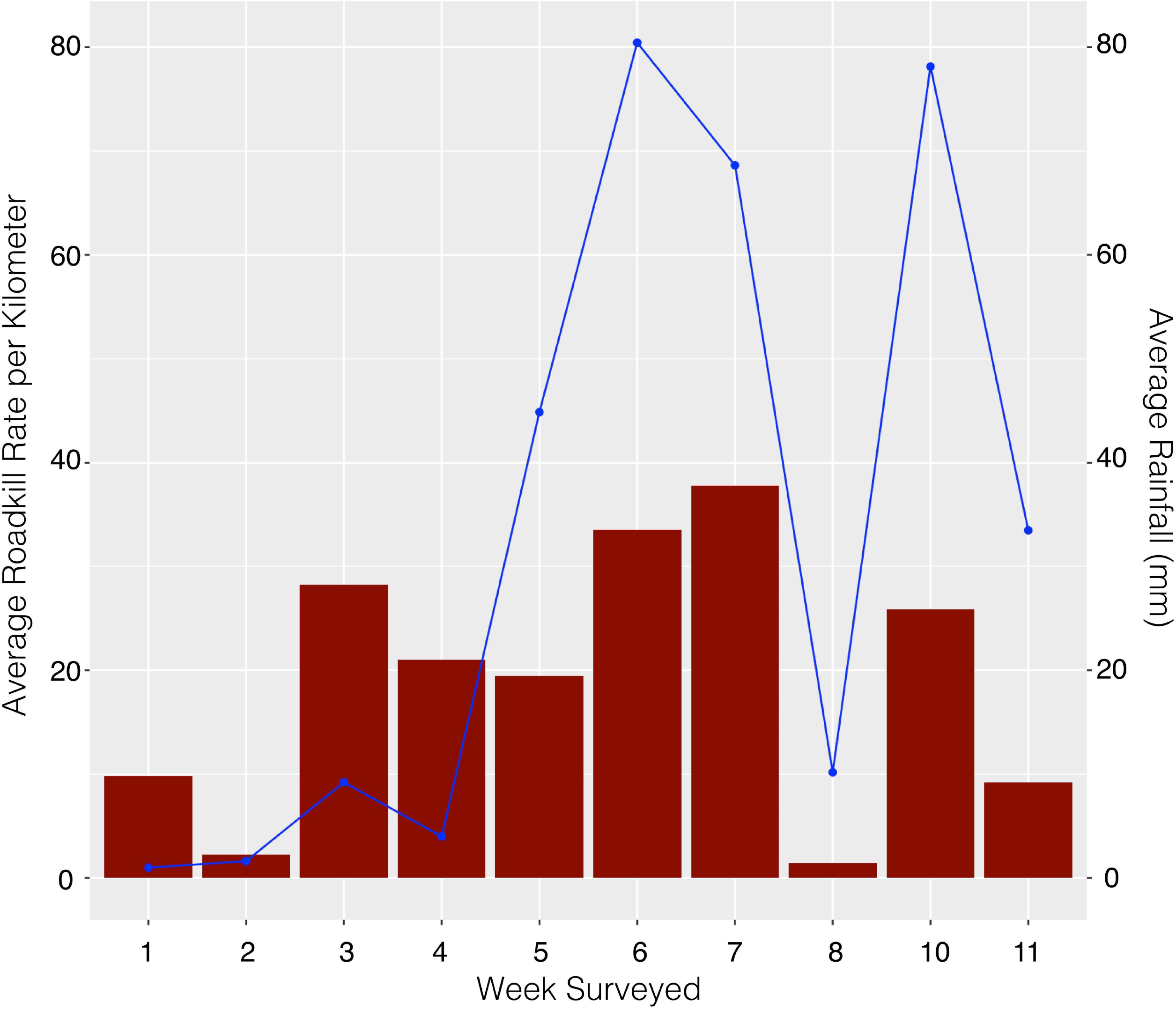

### GLM and predictions

Out of the 15 GLMs, no individual model best explained the variability in MME, with seven models having a ΔAICc of <2 (Table 2). Among the predictor variables considered, none had a 95% beta estimate that did not cross zero, indicating low confidence of any effect or significant impact on the (Supplementary Table 2). In addition, the beta estimates for the traffic volume and the rain are both close to 0, illustrating a negligeable effect on the MME occurrences (Supplementary Table 2).

**Table 2:** GLM results illustrating impact of different variables on the amount of roadkill in HK. The first four models are significantly impacting the mass mortality event occurrences.

|  | **(Intercept)** | **Traffic Volume** | **RH** | **Temp** | **Rain** | **Site** | **df** | **logLik** | **AICc** | **delta** | **weight** |
| --- | --- | --- | --- | --- | --- | --- | --- | --- | --- | --- | --- |
| **model123** | -0.27101 | 0.013671 | 0.115263 | -0.38851 | NA | NA | 4 | -24.8128 | 58.45889 | 0 | 0.172771 |
| **model234** | -0.57399 | NA | 0.112428 | -0.35572 | -0.00133 | NA | 4 | -25.1421 | 59.11758 | 0.658695 | 0.12429 |
| **model25** | -10.9239 | NA | 0.122696 | NA | NA | + | 5 | -23.9517 | 59.18005 | 0.72116 | 0.120469 |
| **model45** | -0.94597 | NA | NA | NA | 0.020582 | + | 5 | -24.0672 | 59.41098 | 0.952091 | 0.107332 |
| **model235** | -0.04372 | NA | 0.090323 | -0.28817 | NA | + | 6 | -23.1442 | 60.11442 | 1.65553 | 0.075505 |
| **model35** | 8.357096 | NA | NA | -0.31067 | NA | + | 5 | -24.4779 | 60.23232 | 1.773432 | 0.071183 |
| **model5** | -0.28768 | NA | NA | NA | NA | + | 4 | -25.767 | 60.36739 | 1.908502 | 0.066534 |
| **model345** | 6.007587 | NA | NA | -0.24661 | 0.017413 | + | 6 | -23.3514 | 60.52881 | 2.069927 | 0.061375 |
| **model245** | -7.23592 | NA | 0.076018 | NA | 0.01145 | + | 6 | -23.6444 | 61.11488 | 2.655994 | 0.045786 |
| **model145** | -0.69888 | -0.0116 | NA | NA | 0.021527 | + | 6 | -23.8931 | 61.61223 | 3.153346 | 0.035705 |
| **model125** | -10.8282 | -0.00086 | 0.121849 | NA | NA | + | 6 | -23.9508 | 61.7276 | 3.268711 | 0.033704 |
| **model124** | -21.0543 | 0.012688 | 0.232087 | NA | -0.00558 | NA | 4 | -26.7877 | 62.40881 | 3.94992 | 0.023975 |
| **model15** | -0.09463 | -0.00796 | NA | NA | NA | + | 5 | -25.6723 | 62.6213 | 4.162409 | 0.021558 |
| **model135** | 8.24808 | -0.0036 | NA | -0.30357 | NA | + | 6 | -24.4595 | 62.74516 | 4.286277 | 0.020264 |
| **model134** | 11.5998 | 0.008818 | NA | -0.46354 | 0.010561 | NA | 4 | -26.9919 | 62.81705 | 4.358167 | 0.019548 |

The averaged model were used to predict the probability of a MME using both the original data and the original data with traffic volume set to a constant mean value. With a prediction probability of > 75% set as a likely MME, both our model-averaged predictions predicted 9 MME, 8 of them being true MME out of 20 (Table 3 and Supplementary Figure 2). Therefore, both models have the same predictive accuracy and include 54% of the total RpK in their prediction (Supplementary Figure 2). Both models also have significant attribution of non-MME with 32 out of 33 non-MME detected (Table 3). The difference in prediction is negligeable between the two models, potentially due to the low impact of traffic volume of the MME occurrence (Supplementary Figure 2).

**Table 3:** Accuracy in MME detection before and after fixing the traffic volume.

| Original model | p > 0.75 | p < 0.75 | Total |
| --- | --- | --- | --- |
| Mass mortality event | 8<br>(True +ve) | 12<br>(False -ve) | 20 |
| No mass mortality event | 1<br>(False +ve) | 32<br>(True -ve) | 33 |
| Total | 9 | 44 | 53 |
| Fixed-Traffic model | p > 0.75 | p < 0.75 | Total |
| Mass mortality event | 8<br>(True +ve) | 12<br>(False -ve) | 20 |
| No mass mortality event | 1<br>(False +ve) | 32<br>(True -ve) | 33 |
| Total | 9 | 44 | 53 |

## Discussion

Amphibian roadkill is a serious threat to biodiversity, with billions of carcasses found on roads each year. By investigating the spatiotemporal distribution of these roadkill for P. *hongkongensis* populations during their migratory season, we confirmed previously observed variations in the timing of mass mortality events and identified sections of road within known hotspots with exceptionally high numbers of roadkill. No external factors significantly explained the timing of mass mortality events in our study, but our current prediction models show accuracy in detection of major mass mortality events (MMEs). By working at a finer spatiotemporal scale, we justify the implementation of site-specific mitigations to protect the newt populations locally and encourage future conservation projects worldwide to consider finer-scale surveys as a better way to understand distribution of roadkill before engaging in any mitigation implementation.

### *P. hongkongensis* roadkill and mass mortality events in Hong Kong

Our results revealed that newts are the primary roadkill victims on the four sites we surveyed (>90% of all carcasses), with over a thousand individuals of this threatened species killed in less than 3 months. These results align with those of previous studies and reinforce the idea that amphibians are the primary victims of roadkill (Beebee, 2013; D’Amigo et al., 2015; Hallisey et al., 2022). We observed contrasting spatial distributions between our survey sites: two having evenly distributed carcasses along the transects (KP and MTL) and the two others having spatially concentrated mortality events (CL and PNS) (Figure 1). This heterogeneity in spatial distribution is characteristic of migratory species (e.g., Clevenger, Chruszcz & Gunson, 2003; Arevalo et al., 2017; Jia et al., 2024). Still, we found most *P. hongkongensis* carcasses in close proximity (<100m) to their core terrestrial habitat and streams in all sites (Figure 1). Those clusters (i.e., hotspots) are mainly found near water bodies (i.e., breeding pools) and high-quality habitats (i.e., feeding areas) such as wetlands intersected by roads (D’Amigo et al., 2015; Zhang et al., 2018; Pinto et al., 2024). The observed peak of mortality in April is similar among our sites. Zhang et al. (2018) described similar results for *R. chensinensis* roadkill distribution, with fewer roadkill in other months. This temporality, with seasonal mortality peaks, corresponds to the typical migration period of amphibians (Arevalo et al., 2017; Canal et al., 2018). In addition, mass mortality events, which are present in a minority of surveys, illustrate those isolated migratory days (Figure 3; Clevenger, Chruszcz, & Gunson, 2003; Garriga et al., 2017). The large number of carcasses found in one day at some survey sites illustrates the significant impact of roads on local newt populations, as it is far greater than previously reported in other amphibian studies (e.g., Hallisey et al., 2022).

### Factors Influencing Mass Mortality Events and Predictions

Rainfall and relative humidity are often considered as main factors triggering roadkill events in amphibians populations (e.g., Zhao et al., 2023; Blais et al., 2024, Pinto et al., 2024). Despite similar apparent patterns between average rainfall and number of carcasses (Figure 4), we did not find a significant relationship between rainfall, relative humidity, and occurrence of mass mortality events (Table 2 and Supplementary Table 2). While rainfall and relative humidity in April may trigger the start of the migratory season in Hong Kong, they do not directly impact mass mortality event occurrences on a daily basis (Parsons, 2021; Jia et al., 2024). Other variables included in our GLM models (temperature, traffic volume, and differences between sites) showed non-significant influence over the probability of MME occurrences (Table 2 and Supplementary Table 2), following results of previous studies (Clevenger, Chruszcz & Gunson, 2003; Hallisey et al., 2022). From our results, we suggest that the significant relationship between similar variables and the mortality events found in other studies might be due to the different scale of surveying the roads (D’Amigo et al., 2015; Morelli et al., 2023; Blais et al., 2024). Unlike most previous studies surveying roads once or twice a month along the entire year, we aimed for weekly to bi-weekly surveys, giving us a finer scale of migratory events and potentially explaining why we do not have strict significance but still observe similarities. Other landscape parameters such as level of urban development or forest composition near the hotspots might be considered as potential factors influencing roadkill but were not included in this study (Sillero et al., 2019; Sousa-Guedes, Franch & Sillero, 2021; Jia et al., 2024).

The development and evaluation of prediction models based on averaged GLMs aimed to provide valuable tools for forecasting future MME events. Due to the non-identification of specific factors triggering MME occurrences through our GLMs, the current predictive power of both models is relatively low (8 out of 20 MMEs detected) (Garriga et al., 2017; Hallisey et al., 2022). However, the models detect the deadliest MMEs, encompassing 54% of total carcasses, thus allowing potential avoidance of the majority of the future roadkill. Future surveys should take into account additional factors that could be significantly triggering MMEs, reinforcing the accuracy of the predicting models (Farmer & Brooks, 2012).

### Implementations of mitigation measures

Of our four study sites, KP should be prioritized due to its large roadkill volume (61% of total carcasses) and because the traffic present at this site is mostly composed of non-residents driving up the hills for a view of the city’s iconic skyline. Both temporary and long-term mitigations could be implemented to safeguard hundreds of newts during their migrations. For example, the government could consider restricting vehicle access to Fei Ngo Shan Road during the peak migration season of *P. hongkongensis* to only residents of local villages or staff of Tate’s Cairn Meterological Station. Alternatively, they could also seasonally convert a downhill portion of Fei Ngo Shan Road to a single lane, two-way road and divert the uphill traffic to Sha Tin Pass Road. The implementations of temporary fences, installation of “newt crossing” road signs, as well as deviating the current road at a nearby road junction (from Fei Ngo Shan road to Jat’s Incline road) during the migratory period of March and April could surely safeguard the vast majority of newts locally (Lesbarreres & Fahrig, 2012; Eberhardt, Mitchell & Fahrig, 2013). Granting permissions to local volunteers to remove this protected species from the roads on specific MME days could also support the conservation efforts, potentially using tailored predictive models to decide when to take action. Despite the fact that newts (and amphibians in general) are migrating in specific seasons, temporary measures alone may not be sufficient to ensure their long-term population persistence (Grilo, Bissonette & Cramer, 2010; Garriga et al., 2017). The implementation of long-term mitigation measures such as culverts and fences can effectively facilitate safe crossing (Dodd, Barichivich & Smith, 2004; Glista, DeVault & DeWoody, 2009; Lesbarreres et al., 2010; Pinto et al., 2024). Overall, mitigation measures for migratory amphibian species are well-documented and highly effective when tailored to the specific groups locally during their peak activity season (D’Amigo et al., 2015).

To ensure effective mitigation measures for amphibians that seasonally migrate, we demonstrated here that preliminary surveys should be conducted to understand the specific local spatiotemporal distribution of roadkill. Site-specific and fine-scaled temporal investigations are necessary because different species in different locations respond differently to weather conditions, which affects the expected roadkill pattern and the predictions of future mass mortality events (Parsons, 2021). Indeed, the non-significance of our predictor variables such as rainfall and relative humidity, at the difference of other studies (e.g., Zhang et al., 2018; Morelli et al., 2023; Blais et al., 2024), shows that a complete understanding of the local amphibian ecology is necessary before development of short- or long-term and potentially costly mitigation measures, to ensure the implementation of the most effective ways to safeguard biodiversity (Farmer and Brooks, 2012). Lastly, our study shows that P. *hongkongensis* requires urgent mitigation measures, continuous monitoring, and a better understanding of the external factors triggering the mass mortality events, in order to improve the prediction models and act more efficiently at the local scale in the future.

## Conclusions

This study sheds light on the spatiotemporal dynamics of roadkill affecting P. *hongkongensis* populations, unveiling previously unexplored variations in mass mortality events and hotspot locations along roads. We offer critical insights into road-related mortality of the Hong Kong newt populations and justify the importance of realising site-specific field surveys before engaging in mitigation measures. In our case, we suggest that refining predictive models and exploring novel variables influencing MME occurrences will be essential for enhancing the effectiveness of mitigation strategies and safeguarding vulnerable amphibian populations in the region. These findings offer valuable insights for amphibian conservation efforts, highlighting the importance of fine-scale surveys and targeted mitigation strategies to effectively safeguard biodiversity in the face of road-related threats.

## Acknowledgements

We thank the backers of the crowdfunding campaign “Life (Cycle) of the Hong Kong Newt” for funding this work. We are grateful to the 20 citizen scientists who collect the roadkill data. We also thank the administrative staff of Lingnan University Science Unit for providing administrative support, and Hazel Chan from Trailwatch, who enabled access to data stored on their platform.

No competing interests between the authors.

## Notes

### Competing Interest Statement

The authors have declared no competing interest.

