## Supplementary figures and images for "Unveiling patterns of roadkill of a migratory amphibian in Hong Kong with implications for mitigation"

### Supplementary_fig1.jpeg

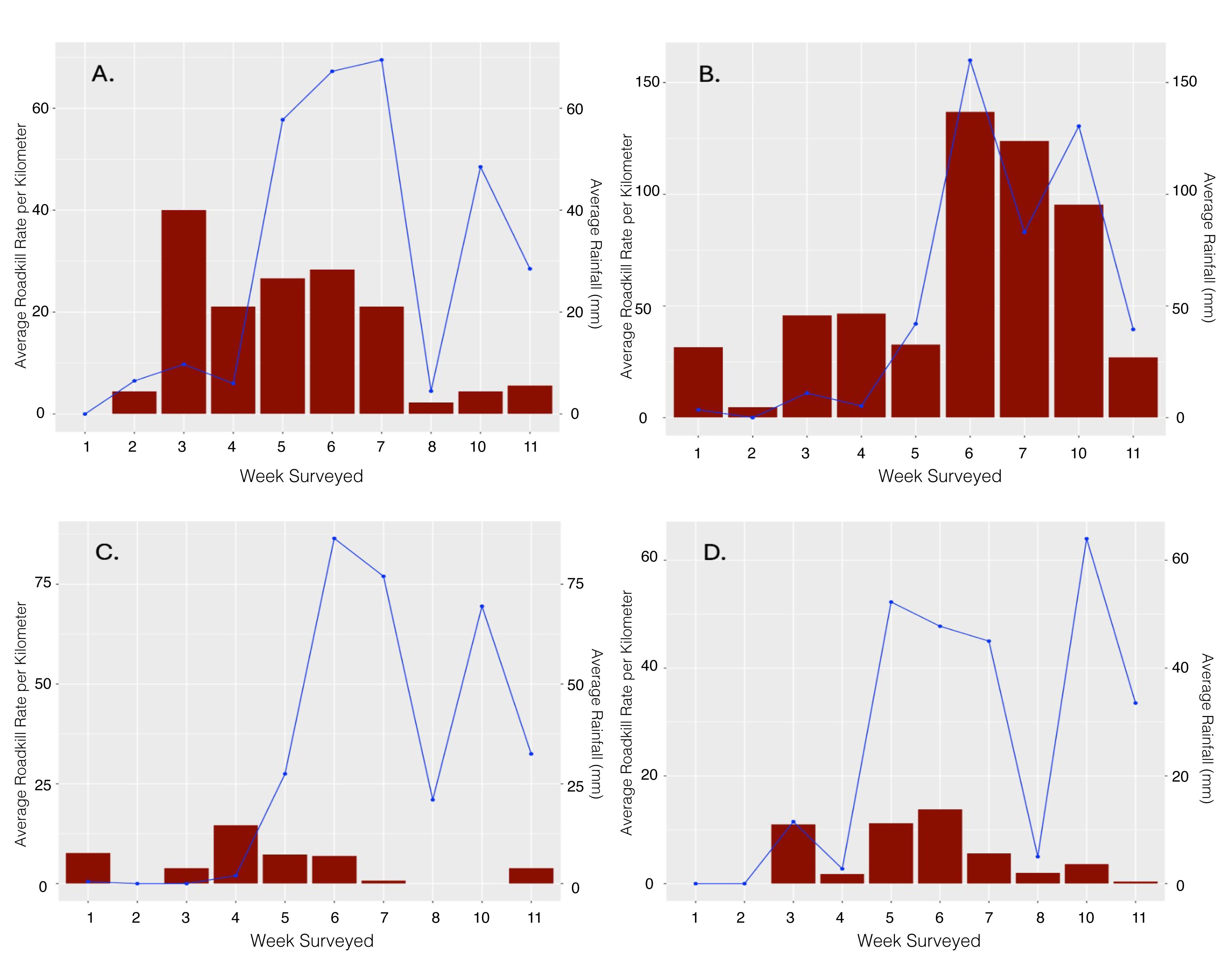

### Supplementary_fig2.png

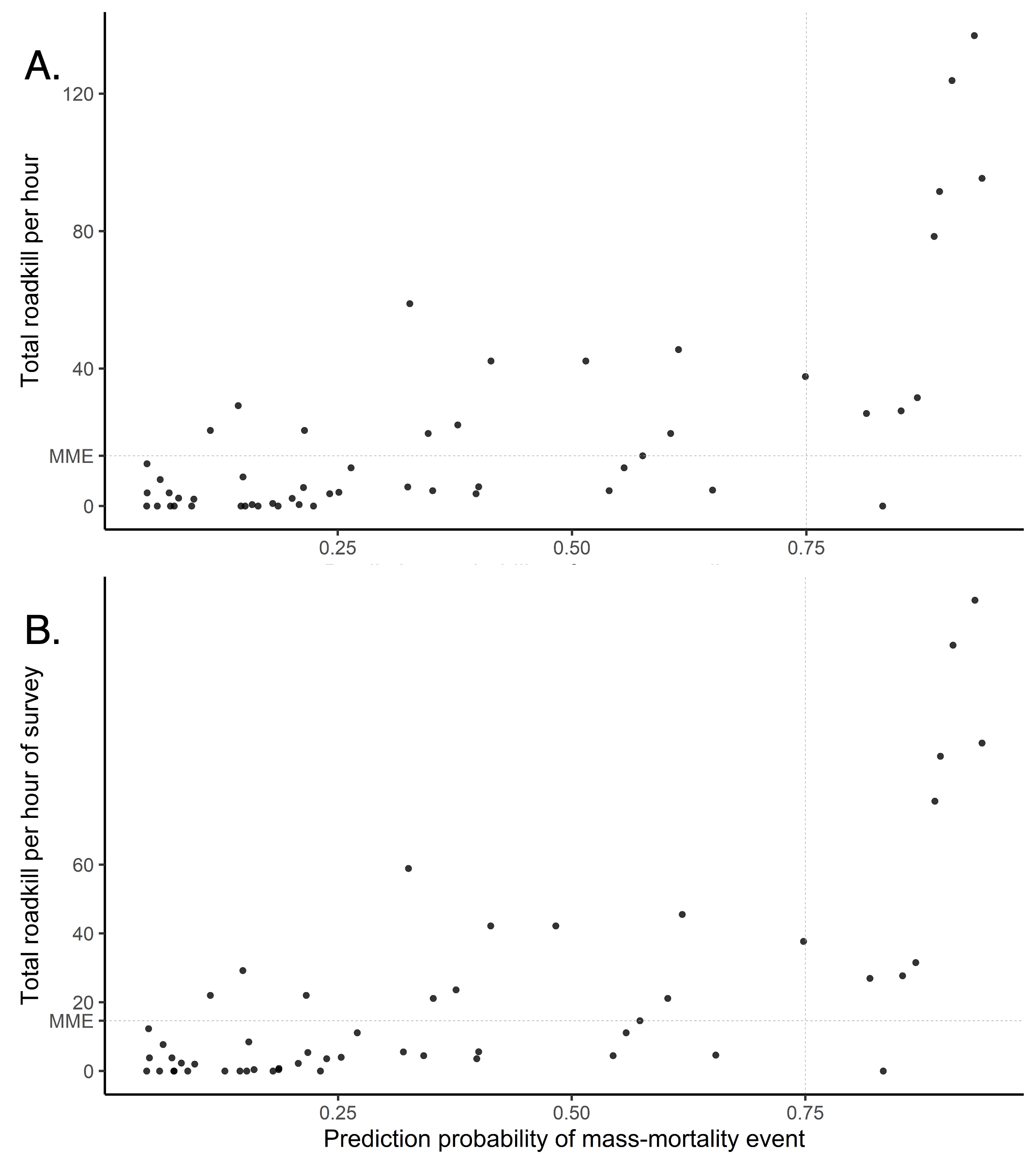
